# Bridging the neural synchronization to linguistic structures and natural speech comprehension

**DOI:** 10.64898/2026.03.23.713668

**Authors:** Jordi Martorell, Giovanni M. Di Liberto, Nicola Molinaro, Lars Meyer

**Affiliations:** Aix-Marseille Université (AMU) & CNRS, Laboratoire Parole et Langage (LPL), Aix-en-Provence, France; Institute for Language, Communication and the Brain (ILCB), Marseille, France; BCBL, Basque Center on Cognition, Brain and Language, Donostia/San Sebastián, Spain; Research Group “Language Cycles”, Max Planck Institute for Human Cognitive and Brain Sciences, Leipzig, Germany; School of Computer Science and Statistics, University of Dublin, Trinity College, Dublin 2, Ireland; ADAPT Centre; Trinity College Institute of Neuroscience and Trinity Centre for Biomedical Engineering, University of Dublin, Trinity College, Dublin 2, Ireland; Ikerbasque, Basque Foundation for Science, Bilbao, Spain; Department of English and Linguistics, Johannes Gutenberg University Mainz, Germany; Clinic for Phoniatrics and Pediatric Audiology, University Hospital Münster, Germany

## Abstract

Speech comprehension involves the inference of abstract information from continuous acoustic signals. Prior work suggests that electrophysiological activity is synchronized with abstract linguistic structures (phrases and sentences) during the processing of isochronous syllable sequences. It is yet unclear whether this prior evidence generalizes to natural speech comprehension, which requires the flexible processing of continuous speech, where syllables and other types of linguistic units are anisochronous. Our magnetoencephalography experiment investigated neural synchronization to acoustic (syllables) and abstract units (phrases and sentences) using continuous speech ranging from artificial isochronous to more natural anisochronous. We find that neural synchronization to phrases and sentences, but not syllables, is resilient to naturalistic anisochrony. This suggests that linguistic structure processing reflects endogenous inferences that are fundamentally distinct from the exogenous processing of syllables driven by speech acoustics. Lateralization and linear regression results extend this functional dissociation as hemispheric asymmetry: stimulus-independent leftward lateralization for linguistic structure processing but stimulus-driven rightward lateralization (or bilaterality) for both syllable and acoustic processing. Our findings provide a more realistic characterization of the flexible neural mechanisms supporting the efficient comprehension of natural speech.

## Introduction

Speech comprehension involves the inference of abstract linguistic units from continuous acoustic signals (Bever and Poeppel, 2010; Martin, 2020, 2016; Meyer et al., 2020a, 2020b). Abstract units come in different durations—smaller units (e.g., words) form larger units (e.g., multi-word phrases). Acoustic cues in speech only allow us to infer the boundaries of some units exogenously (e.g., (Coupé et al., 2019; Ding et al., 2017b; Greenberg, 1999; Inbar et al., 2025, 2020; Rosen, 1992; Zhang et al., 2023)). For instance, only a good half of the multi-word phrases in running speech are accompanied by acoustic boundaries that are strong enough for successful delineation (Giulio Degano et al., 2024). Accordingly, endogenous processes must be involved (Bever and Poeppel, 2010; Halle and Stevens, 1962; Martin, 2020, 2016; Meyer et al., 2020a, 2020b). How do exogenous and endogenous inferences of unit boundaries work together in the brain?

Both exogenous and endogenous types of inference have been linked to narrow-band low-frequency electrophysiological activity (< 8 Hz) recorded with electro– and magnetoencephalography (MEG). Such activity synchronizes with amplitude modulations in the speech envelope (Ahissar et al., 2001; Bourguignon et al., 2013; Luo and Poeppel, 2007). Likewise, neural synchronization with linguistic boundaries occurs without concurrent acoustic modulations, in particular in the delta band (< 4 Hz; for reviews, see (Martorell et al., 2023; Meyer, 2018; Rimmele et al., 2018; Zoefel et al., 2018)).

It is yet unclear whether this prior evidence generalizes to the processing of natural speech due to ecological validity considerations (Alexandrou et al., 2020; Di Liberto and Ip, 2025; Hamilton and Huth, 2018; Nastase et al., 2020). Prior work exclusively used artificial isochronous stimuli in the so-called frequency-tagging paradigm (Buiatti et al., 2009; Nozaradan, 2014; Nozaradan et al., 2011). The approach was pioneered by Ding and colleagues (Ding et al., 2016) who presented isochronous sequences of mono-syllabic words corresponding to 2-syllable phrases (e.g., “new plans”) and 4-syllable sentences (e.g., “new plans gave hope”). As each syllable was adjusted to a duration of 0.25 s, phrases occurred every 0.5 s (i.e., at a frequency of 2 Hz) and sentences every 1 s (i.e., at 1 Hz). While the speech envelopes contained acoustic modulations only at the syllable frequency (i.e., at 4 Hz), MEG revealed neural responses also at the frequency of abstract multi-word structures (i.e., at 2 Hz for phrases and 1 Hz for sentences).

Isochrony falls short of the duration variability of linguistic units in natural speech; moreover, the concatenation of acoustically-independent syllables disregards natural coarticulation. This could lead to a systematic underestimation of neural responses, since isochronous speech impairs word recognition (Aubanel et al., 2016; Aubanel and Schwartz, 2020; Robson et al., 2024) and stimuli containing artificial durations decrease neural synchronization to speech envelope (Chalas et al., 2024; Kayser et al., 2015; Klimovich-Gray et al., 2021). Responses could also be overestimated (Luo and Ding, 2020), as temporal predictability boosts measures of neural synchronization (Haegens and Zion Golumbic, 2018) and removal of coarticulation increases speech-brain synchronization (Deoisres et al., 2023). Concurrently, acoustically-independent isochronous syllables provide a key acoustic cue for synchronization at the syllabic rate. This could also interfere with the processing of multi-word linguistic structures as acoustic isochrony conflates the temporal predictability of linguistic units with and without acoustic correlates. This confound casts doubts on the strictly endogenous nature of multi-word synchronization, echoing discussions about underlying neural mechanisms in other domains (Bouwer et al., 2023, 2020; Breska and Deouell, 2017; Doelling et al., 2019; Morillon et al., 2016).

Our MEG experiment examined whether prior frequency-tagging results generalize to natural speech comprehension. We investigate the impact of isochrony on neural synchronization to linguistic units (syllables, phrases, and sentences) by expanding the frequency-tagging paradigm with anisochronous units that mirror their varying durations that are present in continuous natural speech. We provide evidence that MEG synchronization to abstract phrases and sentences (but not to syllables with acoustic correlates in the stimulus) is resilient to the temporal variability of anisochronous continuous speech. This key finding suggests that the processing of multi-word linguistic structures relies on endogenous inferences that are fundamentally distinct from acoustic-driven exogenous responses, possibly reflecting the flexibility of speech comprehension mechanisms involved in more natural contexts.

## Results

### Continuous anisochronous speech captures neural synchronization to linguistic structures

We aimed to separate the neural synchronization to linguistic structures from the artificial isochrony of frequency-tagging stimuli. Brain activity from 30 healthy participants was recorded non-invasively with MEG while they listened to 20-second speech trials corresponding to 10 consecutive sentences, each composed of two phrases and four bi-syllabic words (Figure 1A). Stimuli were continuous speech streams without acoustic separation between adjacent syllables/words, more similar to natural speech (Figure 1B). In addition, we manipulated the duration of three types of linguistic units (syllables, phrases, and sentences) to create four conditions ranging from artificial isochronous to more natural anisochronous durations in a continuum (Figure 1C). This collaterally manipulated isochrony of bi-syllabic words, which displayed similar duration patterns as syllables (see Figure S1). To control for acoustic modulations, we removed acoustic pitch cues related to the timescales of multi-word linguistic units (see Methods).

**Figure 1.**
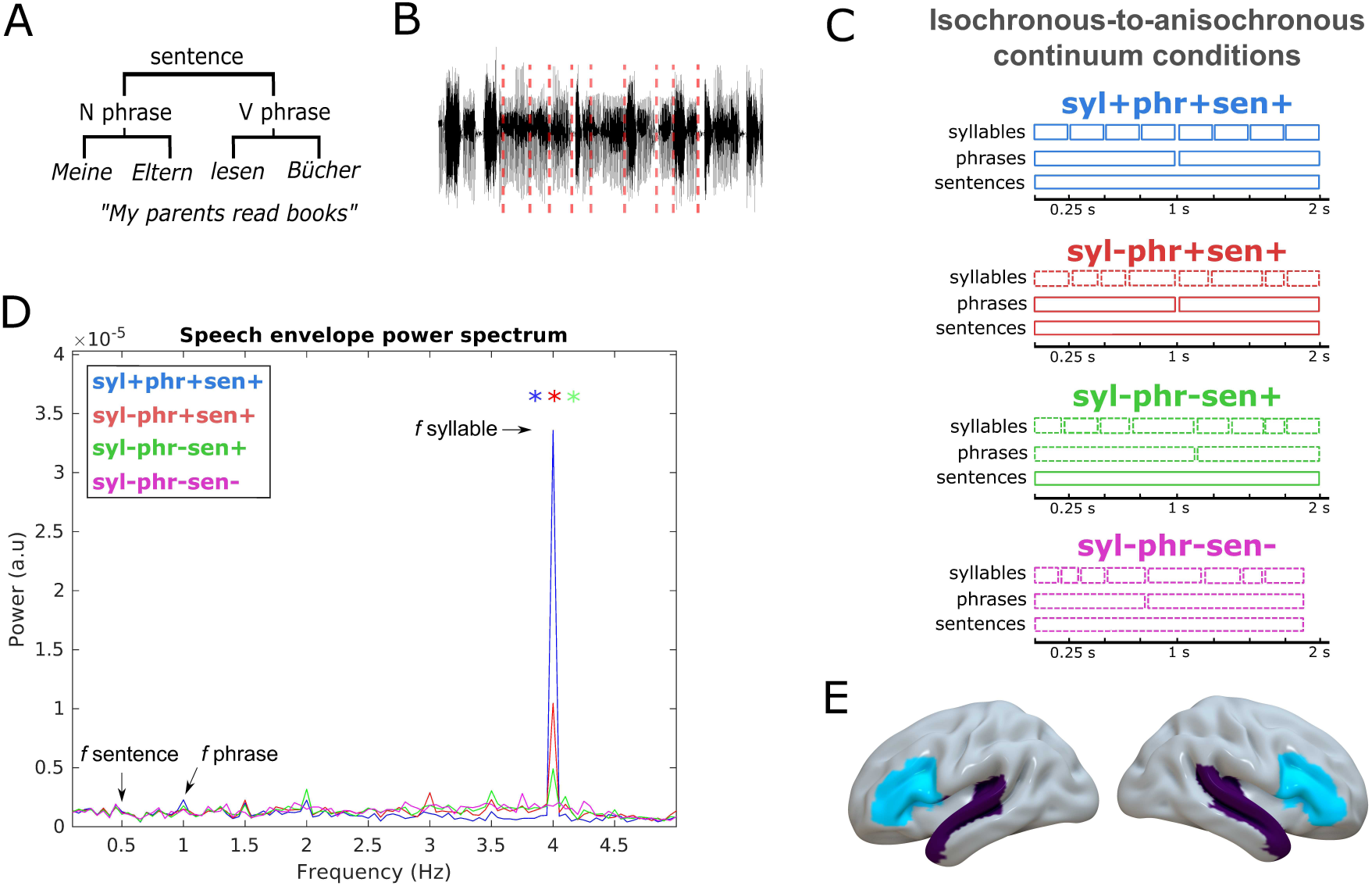
Overview of experimental design. (A) Structure of an example sentence in German, composed of a 2-word Noun phrase and a 2-word Verb phrase (all words are bi-syllabic). (B) Example of synthesized continuous speech trial. Red vertical lines represent the manually annotated syllable boundaries (here shown for one full 8-syllable sentence). (C) Experimental conditions from the current study representing the isochronous-to-anisochronous continuum. Each condition (color-coded) contains a different combination of isochronous (“+”, solid boxes) and/or anisochronous (“-”, dashed boxes) linguistic units (syl = syllables; phr = phrases; sen = sentences). The X-axis represents time (in seconds, s) and displays the different durations of linguistic units (anisochronous durations are for illustrative purposes). Only one sentence is shown but each continuous speech trial contained 10 consecutive sentences. (D) Power spectrum of the speech envelope from each condition (color-coded; see Legend). X-axis represents frequencies and Y-axis power. Arrows indicate frequencies (f) of interest along with their corresponding linguistic unit. Asterisks represent SNR results showing significantly stronger frequency-specific power relative to neighbouring (+/−4) frequency bins (see Methods for further details). (E) Brain regions used for the analyses: Inferior Frontal Gyrus (IFG; cyan) and Superior Temporal Gyrus (STG; purple) from the left and right hemispheres.

We ascertained the efficacy of our acoustic manipulation through spectral analysis (see Methods). As expected, stimuli contained the most salient acoustic modulations at 4 Hz corresponding to the frequency of syllables (Figure 1D). The peak was significantly stronger for isochronous compared to anisochronous syllables (‘syl+phr+sent+’ > ‘syl-phr+sent+’/‘syl-phr-sent+’/‘syl-phr-sent-’) and was reduced across all anisochronous conditions (see Methods). Critically, spectral analysis revealed no differences across conditions at the phrase (1 Hz) and sentence (0.5 Hz) frequencies (see Methods). Accordingly, acoustic cues cannot allow for delineating abstract multi-word linguistic structures (i.e., phrases and sentences) consistently over the isochronous-to-anisochronous continuum.

During each trial of the MEG experiment, participants performed a sentence-matching task. Behavioral accuracy was high (mean (standard deviation); ‘syl+phr+sent+’: 0.7 (0.15); ‘syl-phr+sent+’: 0.655 (0.13); ‘syl-phr-sent+’: 0.685 (0.13); ‘syl-phr-sent-’: 0.71 (0.1)) and comparable across conditions (condition effect: F(3,116) = 1.1191, p > .05). Behavioral results thus suggest that participants were paying attention to our continuous stimuli similarly across conditions comprising isochronous-to-anisochronous speech.

To confirm the methodological validity of our continuous speech paradigm, we first assessed MEG synchronization to linguistic units through Inter-Trial Phase Coherence (ITPC). ITPC is commonly used in frequency-tagging studies due to its high reliability to detect periodic patterns in signals (see Methods). Analyses were conducted in MEG source space, where we estimated single-trial activity from 162 sub-regions (Human Brainnetome Atlas (Fan et al., 2016); see Methods). We focused on bilateral Inferior Frontal Gyrus (IFG) and Superior Temporal Gyrus (STG; see Figure 1E), as these brain regions are typically involved in both endogenous and exogenous aspects of language processing (Friederici, 2011; Molinaro and Lizarazu, 2018; Park et al., 2015). ITPC was computed in terms of non-uniform distribution of phase angles across trials as expressed by the Rayleigh test’s z-values (see Methods), which were then averaged across IFG/STG regions and left/right hemispheres to reduce the dimensionality of this confirmatory analysis. Statistical significance was quantified by comparing z-values at each frequency of interest with the average of its (+/− 4) neighbouring frequencies (i.e., +/− 0.22 Hz; one-sided *t*-test, uncorrected) across conditions (see Methods).

Significant ITPC was observed at the frequency of each linguistic unit when all units were isochronous (Figure 2). Specifically, in the ‘syl+phr+sent+’ condition, we observed significant ITPC at the syllable frequency (4 Hz; t(58) = 13.426, p < .001), phrases (1 Hz; t(58) = 11.084, p < .001), and sentences (0.5 Hz; t(58) = 8.460, p <.001). This confirms the methodological validity of our paradigm and extends prior frequency-tagging results, as our stimuli involve continuous speech without artificial gaps between syllables. ITPC peaked at the word frequency (2 Hz; t(58) = 12.393, p < .001), possibly related to the behavioural relevance of words to perform the sentence-matching task. ITPC was also significant (p < .05) at the frequency of most linguistic units for the rest of conditions containing anisochronous units. Specifically, in the ‘syl-phr+sent+’ and ‘syl-phr-sent+’ conditions, ITPC was significant at the frequency of each linguistic unit (p < .05). In the ‘syl-phr-sent-’ condition in which all units were anisochronous, we instead observed significant phase synchronization only at the frequency of phrases and sentences (but neither words nor syllables; p > .05). Isochrony is thus not strictly necessary to observe phase synchronization at the frequency of linguistic units, especially for larger units such as phrases and sentences which can elicit reliable frequency-tagged neural responses also with more natural anisochronous speech.

**Figure 2.**
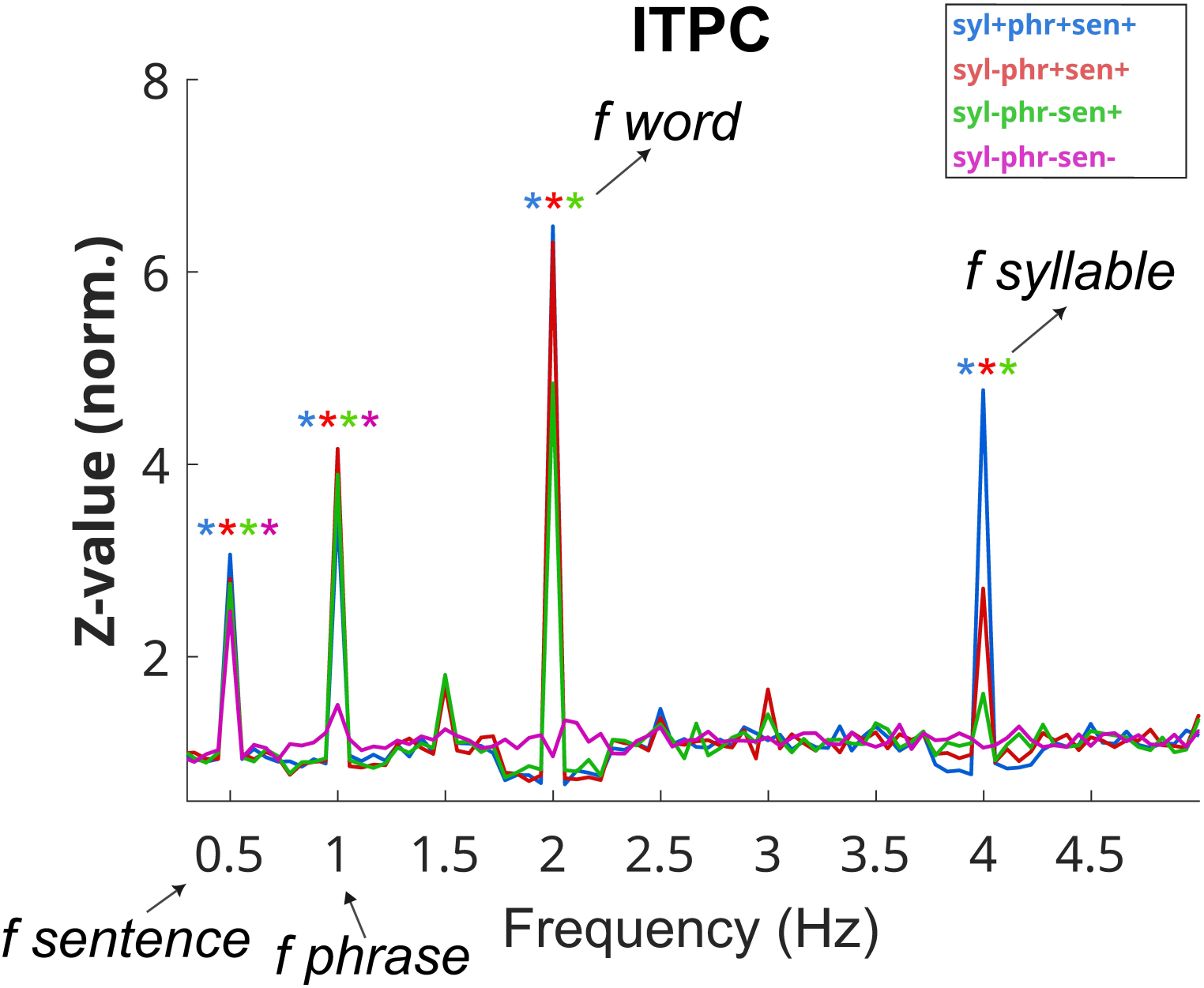
Inter-Trial Phase Coherence (ITPC) spectrum. ITPC averaged across regions (IFG/STG) and hemispheres (left/right), shown separately for each condition across the isochronous-to-anisochronous continuum (color-coded; see Legend). X-axis represents frequencies and Y-axis z-values (normalized). Arrows indicate frequencies (f) of interest along with their corresponding linguistic unit (here including the 2-Hz word frequency). Asterisks represent SNR results showing significantly stronger frequency-specific ITPC relative to neighbouring (+/−4) frequency bins (see also Methods and Figure S2)

Finally, we assessed between-condition ITPC differences only at the syllable frequency (4 Hz), where also spectral analysis of the speech envelope had revealed the most salient acoustic modulations. The 4-Hz ITPC peak was strongest with isochronous syllables and mirrored the decrease across conditions with additional anisochronous units from the spectral analysis of the speech envelope (see Figure S2). Together, isochronous syllables maximize ITPC at the syllable frequency, possibly following the acoustic modulations from the speech envelope.

### Isochrony drives neural synchronization to syllables but not multi-word structures

We then investigated how isochrony impacts MEG synchronization with linguistic structures by computing inter-event phase coherence (IEPC). Unlike ITPC, IEPC examines frequency-specific instantaneous phase synchronization across event boundaries corresponding to the exact occurrence of each linguistic unit (i.e., offset of the preceding unit and onset of the following unit) in the continuous speech signal —thus providing a more precise estimation of neural synchronization to anisochronous units. We hypothesized that anisochronous units would generally decrease IEPC. Crucially, as our conditions progressively contained fewer isochronous units, we were able to assess the selective impact of isochrony for each linguistic unit. IEPC was computed using the instantaneous phase from source-localized single trials and then implementing the non-uniformity’s Rayleigh test (see Methods). Statistical significance was quantified by comparing (normalized) z-values across conditions, IFG/STG regions, and left/right hemispheres separately for each linguistic unit (linear model and bonferroni-corrected *t*-test for post-hoc contrasts; see Methods).

IEPC at syllable boundaries differed across conditions (condition effect: F(3,464) = 111.950, p < .001; Figure 3A and Figure S3A). As expected, IEPC increased for isochronous compared to anisochronous syllables (‘syl+phr+sent+’ vs. ‘syl-phr+sent+’: t(464) = 14.255, p < .001; ‘syl+phr+sent+’ vs. ‘syl-phr-sent+’: t(464) = 14.854, p < .001; ‘syl+phr+sent+’ vs. ‘syl-phr-sent-’: t(464) = 15.649, p < .001). However, in contrast to the speech envelope power and ITPC spectra, IEPC did not differ between conditions with additional anisochronous units (‘syl-phr+sent+’ vs. ‘syl-phr-sent+’: t(464) = 0.599, p > .05; ‘syl-phr+sent+’ vs. ‘syl-phr-sent-’: t(464) = 1.394, p > .05; ‘syl-phr-sent+’ vs. ‘syl-phr-sent-’: t(464) = 0.795, p > .05). We also found that isochrony was associated with right-hemispheric lateralization of responses in general and in the STG in particular (region effect: F(1,464) = 32.378, p < .001; hemisphere effect: F(1,464) = 45.937, p < .001; region x hemisphere interaction: F(1,464) = 2.160, p > .05; condition x region interaction: F(3,464) = 9.1098, p < .001; condition x hemisphere interaction: F(3,464) = 10.901, p < .001; condition x hemisphere x region interaction: F(3,464) = 0.804, p > .05), as it was only observed with isochronous syllables (p < .001) but not with anisochronous syllables (p > .05). Together, syllable-level IEPC appears generally stronger in the right STG, although artificial isochronous syllables particularly maximize this right-STG dominance.

**Figure 3.**
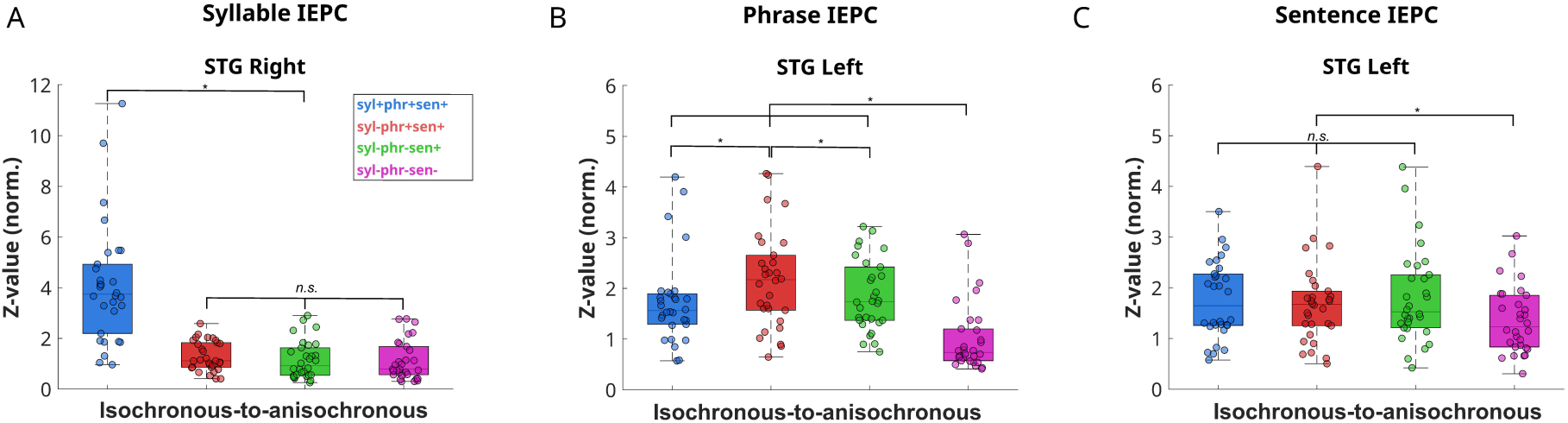
Inter-Event Phase Coherence (IEPC) main results. IEPC derived from instantaneous phase synchronization at frequency-specific linguistic boundaries for syllables (A), phrases (B), and sentences (C), shown separately for each condition across the isochronous-to-anisochronous continuum (color-coded; see Legend) in boxplots (box central line = median; box outlines = 1st and 3rd quartiles; whiskers = 1.5 interquartile range; dots represent single-participant values), only for the region with the strongest effect (regions with less strong effects in Figure S3). X-axis represents conditions and Y-axis IEPC z-values (normalized). Asterisks represent statistically significant differences between conditions (see Methods).

IEPC at phrase boundaries also differed across conditions (condition effect: F(3,464) = 34.253, p < .001; Figure 3B and Figure S3B). Isochronous phrases elicited higher IEPC with anisochronous syllables compared to isochronous syllables (‘syl+phr+sent+’ vs. ‘syl-phr+sent+’: t(464) = −4.158, p < .001). This enhancement effect for anisochronous syllables was reduced under the additional presence of anisochronous phrases (‘syl-phr+sent+’ vs. ‘syl-phr-sent+’: t(464) = 3.474, p < .01), making phrase-level IEPC comparable to the condition comprising only isochronous units (‘syl+phr+sent+’ vs. ‘syl-phr-sent+’: t(464) = −0.684, p > .05). This means that anisochronous syllables enhance phrase-level IEPC, although this boosting effect is neutralized by the lower temporal regularity of anisochronous phrases. The condition additionally containing anisochronous sentences showed the lowest phrase-level IEPC compared to the rest of conditions (‘syl+phr+sent+’ vs. ‘syl-phr-sent-’: t(464) = 5.819, p < .001; ‘syl-phr+sent+’ vs. ‘syl-phr-sent-’: t(464) = 9.977, p < .001; ‘syl-phr-sent+’ vs. ‘syl-phr-sent-’: t(464) = 6.503, p < .001), possibly related to the lowest temporal regularity of phrases when all linguistic units were anisochronous. We also observed that phrase-level IEPC was strongest in the left STG (region effect: F(1,464) = 16.120, p < .001; hemisphere effect: F(1,464) = 53.158, p < .001; region x hemisphere interaction: F(1,464) = 6.433, p < .05) as corroborated by the selective effect in the left (t(464) = −4.632, p < .001) but not right (t(464) = −1.045, p > .05) STG similarly across conditions (condition x region interaction: F(3,464) = 0.976, p > .05; condition x hemisphere interaction: F(3,464) = 1.007, p > .05; condition x region x hemisphere interaction: F(3,464) = 1.062, p > .05). This pattern of results suggest that anisochronous syllables enhance phrase-level IEPC, which is nevertheless robustly strongest in the left STG across the isochronous-to-anisochronous continuum.

IEPC at sentence boundaries was also significantly different across conditions (condition effect: F(3,464) = 8.836, p < .001; Figure 3C and Figure S3C). As expected, we found significantly higher IEPC for isochronous compared to anisochronous sentences (‘syl+phr+sent+’ vs. ‘syl-phr-sent-’: t(464) = 4.790, p < .001; ‘syl-phr+sent+’ vs ‘syl-phr-sent-’: t(464) = 3.423, p < .01; ‘syl-phr-sent+’ vs. ‘syl-phr-sent-’: t(464) = 3.936, p < .001). In addition, we observed comparable sentence-level IEPC across conditions containing isochronous sentences regardless of the isochrony of syllables or phrases (‘syl+phr+sent+’ vs. ‘syl-phr+sent+’: t(464) = 1.367, p > .05; ‘syl+phr+sent+’ vs. ‘syl-phr-sent+’: t(464) = 0.854, p > .05; ‘syl-phr+sent+’ vs. ‘syl-phr-sent+’: t(464) = −0.514, p > .05). These results suggest that, unlike phrase-level effects, sentence-level IEPC is insensitive to the presence of more artificial or natural durations from the smaller linguistic units that concurrently occur in the stimulus. Similarly to phrase-level effects, we also observed that sentence-level IEPC was selectively strongest in the left STG (region effect: F(1,464) = 28.324, p < .001; hemisphere effect: F(1,464) = 54.950, p < .001; region x hemisphere interaction: F(1,464) = 6.433, p < .05), again corroborated by the significant effect in the left (t(464) = −5.373, p < .001) but not right (t(464) = −2.154, p > .05) STG consistently across conditions (condition x region: F(3,464) = 0.683, p > .05; condition x hemisphere: F(3,464) = 0.2042, p > .05; condition x region x hemisphere: F(3,464) = 0.0374, p > .05).

To summarize, we found that isochrony enhances MEG synchronization to syllables, unlike phrase- and sentence-level effects which were instead reduced or unaffected. Moreover, we also report that isochrony maximized the right-STG dominance of syllable-level synchronization but did not alter the left-STG dominance for phrases and sentences. These results clearly suggest that the processing of multi-word structures (but not smaller units such as syllables) flexibly adapts to the temporal variability of anisochronous speech as required to efficiently comprehend natural speech. The word-level IEPC results (Figure S4) support this possibility, showing strong effects of isochrony with consistent bilateral patterns.

### Hemispheric asymmetry for multi-word and syllable processing

We further investigated how isochronous-to-anisochronous speech modulated the hemispheric asymmetry across linguistic units suggested by the IEPC magnitude effects. To this end, we computed a lateralization index (LI; see Methods) on the IEPC values and statistically compared the LI outcome against the empirical bilateral distribution (t-test, two-tailed, corrected for multiple comparisons; see Methods).

For syllables, lateralization varied depending on isochrony (Figure 4A). The most reliable right-hemisphere lateralization emerged with isochronous syllables in both IFG and STG. The presence of anisochronous syllables instead resulted in more bilateral patterns across regions; bilaterality was less pronounced when all linguistic units were anisochronous (i.e., rightward lateralization was not significant after correction for multiple comparisons). These LI patterns extend the previous IEPC results by suggesting that artificial isochronous syllables drive the right-hemisphere lateralization of syllable-level IEPC not only in STG but also in IFG.

**Figure 4.**
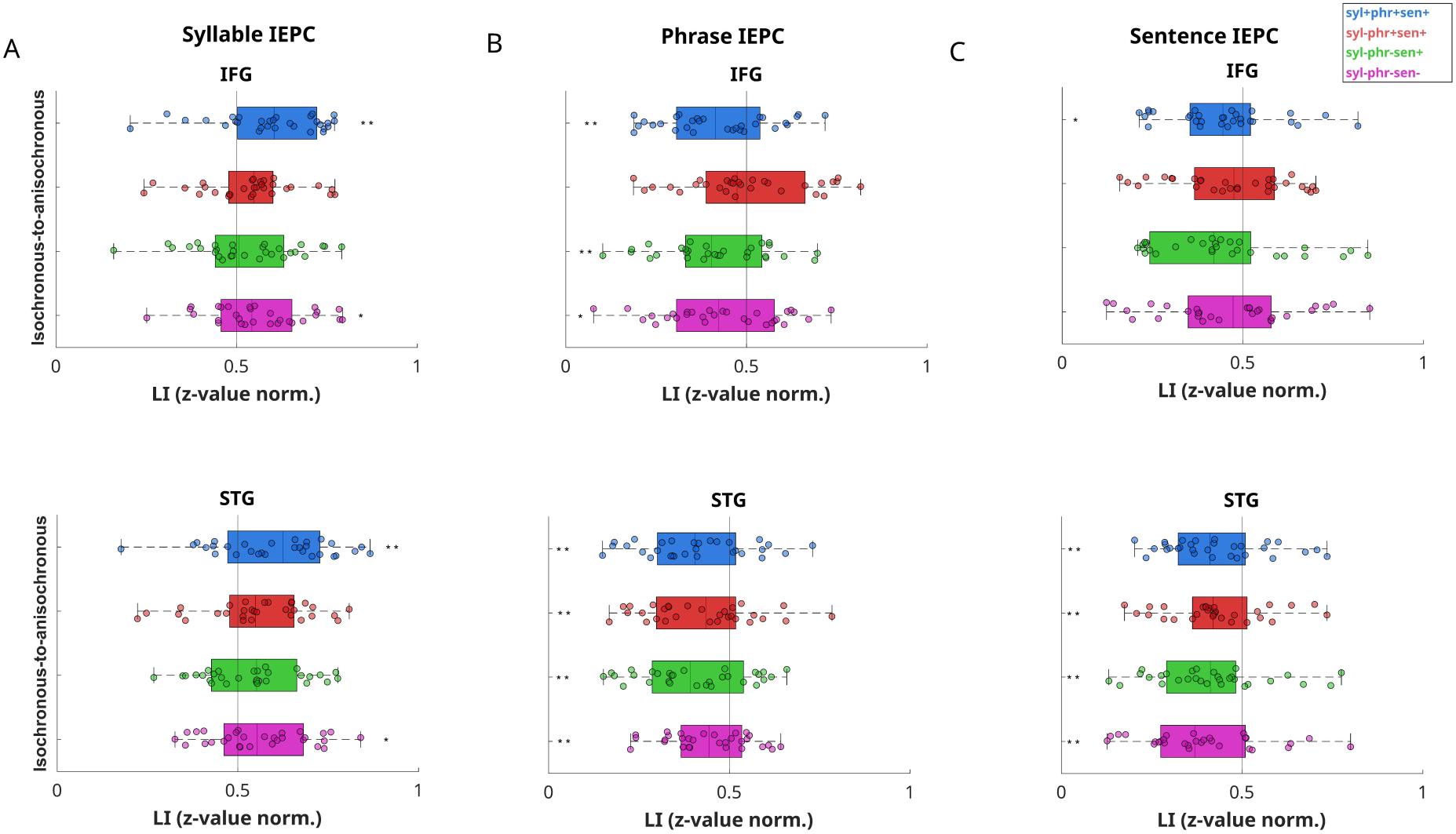
Lateralization Index (LI) of Inter-Event Phase Coherence (IEPC) results. Hemispheric lateralization patterns for syllables (A), phrases (B), and sentences (C) in IFG (upper) and STG (lower), shown separately for each condition across the isochronous-to-anisochronous continuum (color-coded; see Legend) in boxplots (box central line = median; box outlines = 1st and 3rd quartiles; whiskers = 1.5 interquartile range; dots represent single-participant averaged values). X-axis represents LI values (0 = leftward lateralization; 0.5 = bilaterality, marked by vertical dashed line; 1 = rightward lateralization) and Y-axis conditions. Asterisks represent statistically significant differences between conditions before (*) or after (**) correction for multiple comparisons (see Methods and Figure S5).

For phrases, we found distinct IEPC lateralization patterns between IFG and STG for certain conditions (Figure 4B). In IFG, leftward lateralization was significant when all linguistic units were isochronous, but displayed a bilateral distribution when only syllables were anisochronous. The additional presence of anisochronous phrases again resulted in a leftward lateralization, which was less pronounced when all linguistic units were anisochronous (non-significant after correction for multiple comparisons). Considering the similar condition patterns observed in the main IEPC analysis, these results suggest that the enhanced phrase-level IEPC effects for anisochronous syllables corresponded to a relative increase in the right IFG that shifted the overall IEPC towards a bilateral IFG distribution. In contrast, STG consistently displayed significant leftward lateralization across all conditions. These results underline the left-STG dominance for processing phrases across the isochronous-to-anisochronous continuum as previously suggested by the IEPC findings.

For sentences, we again observed distinct IEPC lateralization patterns in IFG and STG (Figure 4C). In IFG, we only found evidence for leftward lateralization when all linguistic units were isochronous, although this asymmetry was not significant after correction for multiple comparisons. In STG, the LI results instead indicated a consistent leftward lateralization across all conditions. Similarly to the STG hemispheric asymmetry for phrases, these findings strengthen the interpretation of the previous sentence-level IEPC results about the insensitivity of the left-STG dominance to isochrony.

Taken together, the LI results corroborate the IEPC hemispheric asymmetry showing that isochrony drives rightward lateralization for syllables independently of the left-STG dominance for phrases and sentences. The word-level LI results (Figure S5) confirm the bilateral distribution across the isochronous-to-anisochronous continuum.

### STG responses to speech acoustics are insensitive to isochrony

Next, we examined how isochrony modulates the processing of speech acoustics in the time domain. Although the speech envelope contained the most salient acoustic modulations at the frequency of syllables, IEPC dynamics at syllable boundaries might not necessarily provide evidence for acoustic-level processing. This is because abstract linguistic units such as syllables do not always coincide with acoustic boundaries (Coupé et al., 2019; Ding et al., 2017b; Zhang et al., 2023). In addition, syllable-level and acoustic-level processing produces similar but yet dissociable neural responses (Brodbeck et al., 2018; Schmidt et al., 2023). We therefore assessed the time-resolved linear relationship between neural responses and the continuous speech envelope of our stimuli using a forward Temporal Response Function (TRF) approach (Crosse et al., 2021, 2016). Specifically, following a lagged ridge regression procedure, we model the contribution of the speech envelope to predict MEG responses in bilateral STG and evaluate the model’s prediction accuracy by means of Pearson’s correlation between predicted and empirical neural data with cross validation (see Methods for further details).

We observed similar prediction accuracy across conditions (condition effect: F(3,232) = 1.349, p > .05; Figure 5). In addition, we found stronger prediction accuracy in the right compared to the left STG (hemisphere effect: F(1,232) = 6.068, p < .05). This right-hemisphere dominance aligns with the observed IEPC effects at syllable boundaries, although in this case such a hemispheric asymmetry was not different across conditions (condition x hemisphere interaction: F(3,232) = 0.089, p > .05). The similarity of prediction accuracy across isochronous-to-anisochronous speech is in line with the notion that speech acoustics are processed in a context-invariant manner.

**Figure 5.**
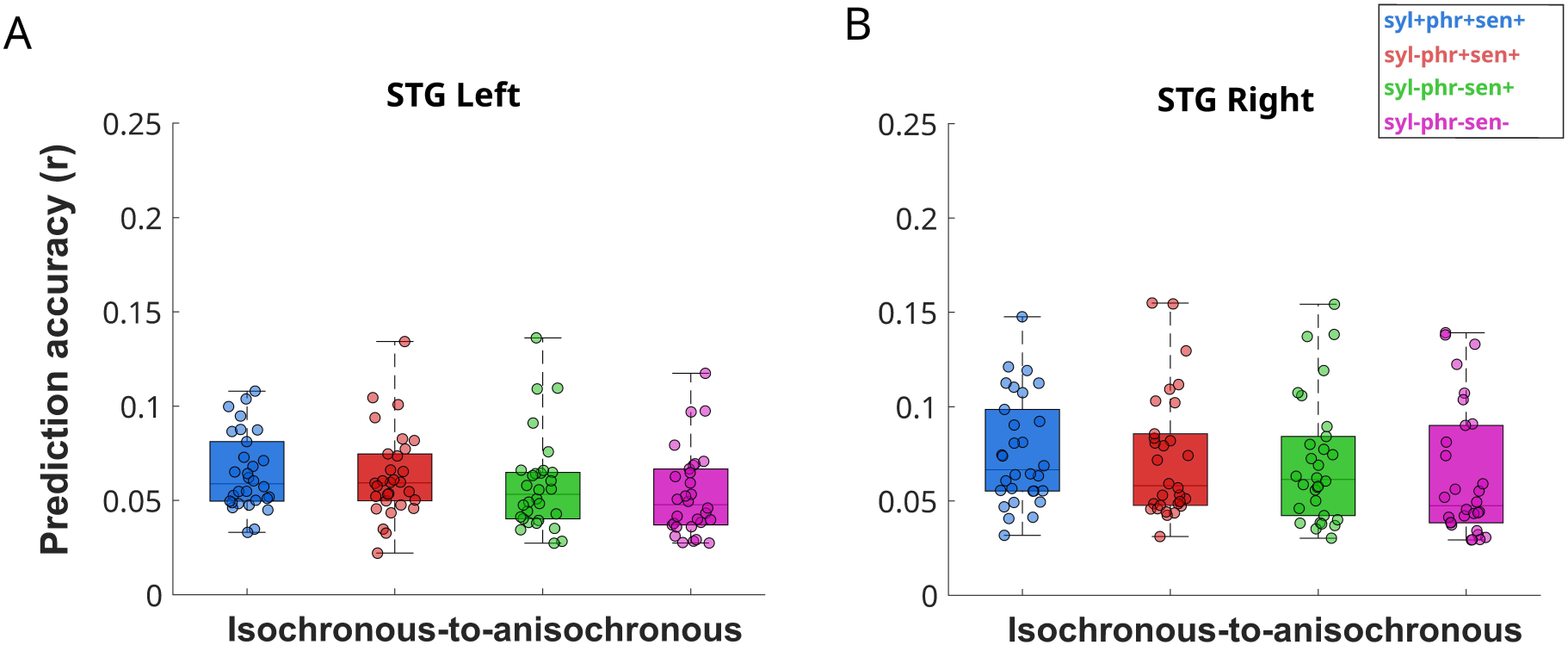
Temporal Response Function (TRF) results. Forward TRF modelling of the speech envelope regressor for left STG (A) and right STG (B), shown separately for each condition across the isochronous-to-anisochronous continuum (color-coded; see Legend) in boxplots (box central line = median; box outlines = 1st and 3rd quartiles; whiskers = 1.5 interquartile range; dots represent single-participant averaged values). X-axis represents conditions and Y-axis the TRF prediction accuracy (Pearson’s r) between predicted and empirical MEG data. No statistical difference between conditions was observed, only a hemisphere effect (right STG > left STG; see also Figure S6).

Unlike TRF results, syllable-level IEPC (Figure 3A) revealed neural enhancement for isochronous syllables maximized in the right STG. To better understand these contrasting results, we computed the correlation between the syllable-level IEPC and the acoustic-level TRF prediction accuracy at right STG (see Methods for further details). The IEPC-TRF correlation results (Figure S6) reflected a positive linear relationship between both measures in all conditions except for the condition containing isochronous syllables. Interestingly, the correlation coefficient was progressively larger across conditions with additional anisochronous units (i.e., ‘syl+phr+sent+’ > ‘syl-phr+sent+’ > ‘syl-phr-sent+’ > ‘syl-phr-sent-’). These results suggest that artificial isochronous syllables give rise to maximized syllable-level IEPC patterns that dramatically differ from the acoustic-level responses observed during the processing of more natural anisochronous speech.

## Discussion

Neural synchronization to multi-word linguistic structures (phrases and especially sentences), but not to syllables, is resilient to anisochrony of continuous speech signals akin to natural speech. Our MEG results thus suggest that the processing of abstract linguistic structures relies on endogenous inferences that are fundamentally distinct from the exogenous processing of syllables driven by speech acoustics. This has implications for the characterization of the neural mechanisms supporting natural speech comprehension.

Neural synchronization to abstract multi-word structures was not driven by artificial isochronous speech. For phrases, we observed enhanced IEPC with more natural anisochronous syllables, although this boosting effect was reduced with the additional presence of anisochronous phrases (Figure 3B). Such stimulus-sensitive modulations are compatible with the compensatory trade-off dynamics between exogenous and endogenous responses observed during more natural anisochronous speech comprehension (Bai et al., 2022; Klimovich-Gray et al., 2021; Meyer et al., 2016; Molinaro et al., 2021; Pérez-Navarro et al., 2024; Reetzke et al., 2021; Tezcan et al., 2023; Zou et al., 2019); see (Meyer et al., 2020a, 2020b). For sentences, neural synchronization was robustly resilient to the temporal variability of anisochronous speech, showing comparable IEPC regardless of the isochrony of syllables and/or phrases (Figure 3C). This suggests that the processing of abstract sentences involves endogenous responses that are independent from stimulus features. Together, our evidence clearly supports that the processing of abstract multi-word structures relies on endogenous inferences that cannot be accounted for by the exogenous processing of speech acoustics. This possibly reflects the flexibility of speech comprehension mechanisms to efficiently adapt to the anisochrony of natural speech.

In contrast to multi-word structures, neural synchronization to syllables was strongly dependent on isochrony (Figure 3A). These patterns are in line with recent findings (López-Madrona et al., 2025; Oganian et al., 2023; Wang et al., 2025; Zou et al., 2021) relating the processing of syllables (and generally linguistic units with acoustic correlates) to exogenous, stimulus-driven responses time-locked to speech acoustics. More concretely, the instantaneous phase of such stimulus-driven responses would exhibit high consistency at regular (i.e., isochronous) boundaries but random fluctuations at variable (i.e., anisochronous) boundaries. Considering the correspondence at the 4-Hz syllable frequency between the speech envelope (Figure 1D) and ITPC (Figure 2 and Figure S2) spectra, the similar TRF prediction accuracy (Figure 5) across the isochronous-to-anisochronous continuum also reinforces the involvement of stimulus-driven responses following the acoustic modulations contained in the speech envelope. Apart from supporting the separation between acoustic-level and syllable-level responses (Brodbeck et al., 2018; Schmidt et al., 2023), the similarity between acoustic-level TRF and syllable-level IEPC except for isochronous syllables (Figure S6) also raises concerns about the ecological validity of isochronous speech to motivate accounts of syllable processing (Assaneo et al., 2019). Our findings thus converge on the stimulus-driven nature of syllable processing, which is likely dramatically maximized with artificial isochronous speech.

This functional dissociation between multi-word structure and syllable processing is further confirmed by the hemispheric asymmetry findings. For both phrase- and sentence-level synchronization, we observed leftward lateralization in STG consistently across the isochronous-to-anisochronous continuum (see Figure 4B and 4C). The selectivity of endogenous responses in temporal (rather than frontal) regions for multi-word processing aligns with recent proposals(Matchin and Hickok, 2019) and departs from more classical views (Friederici, 2011; Jakuszeit et al., 2013). Interestingly, frontal (but not temporal) regions displayed a relative sensitivity to isochrony (especially for phrases), similarly to the endogenous modulations observed during acoustic-level processing (Lamekina et al., 2024; Molinaro and Lizarazu, 2018; Park et al., 2015). Syllable-level synchronization was instead rightward lateralized with isochronous syllables or bilateral with anisochronous syllables (see Figure 4A). Together with this selective effect of isochrony, the rightward lateralization of acoustic-level TRF results strengthen the link between stimulus-driven responses and syllable processing. Importantly, the reported functional and hemispheric dissociation between multi-word structure and syllable processing bridge previous findings from both isochronous (e.g., (Ding et al., 2017a; Glushko et al., 2022; Gui et al., 2020; Lu et al., 2023; Makov et al., 2017; Sokoliuk et al., 2021)) and anisochronous (Broderick et al., 2022; Lizarazu et al., 2019; Molinaro and Lizarazu, 2018; Pefkou et al., 2017) speech comprehension paradigms.

Our findings can also inform ongoing debates about neural mechanisms distinguishing between transient evoked and sustained oscillatory responses (e.g., (Lalor and Nidiffer, 2025; Meyer et al., 2020a; Molinaro, 2025; Oganian et al., 2023; Zoefel et al., 2018; Zou et al., 2021)). Indeed, the stimulus-driven characteristics of syllable-level findings across isochronous-to-anisochronous speech seem to provide reliable evidence for evoked responses - although such effects could additionally reflect oscillatory responses phase resetted by acoustic cues (Doelling and Assaneo, 2021; Giraud and Poeppel, 2012). However, even with anisochronous speech, long-lasting evoked responses at phrase/sentence boundaries (e.g., endogenous language-related components such as the P600) can theoretically result in neural dynamics indistinguishable from sustained oscillatory responses (at least if they tolerate certain deviations from perfect isochrony; see (Doelling and Assaneo, 2021; Rimmele et al., 2018); cf. (Bouwer et al., 2023; Breska and Deouell, 2017)). Importantly, it is also conceivable that evoked and oscillatory responses coexist during speech comprehension (Forseth et al., 2020), possibly reflecting mutually dependent aspects of the same underlying neural mechanisms (Henke and Meyer, 2021; Meyer, 2018); see also (Klimesch et al., 2007; Sauseng et al., 2007)). We thus suggest that characterizing speech comprehension mechanisms might benefit from clarifying the relative contribution of multiple components (e.g., endogenous/exogenous, transient/sustained, etc.) beyond the evoked-oscillatory dichotomy (for further discussion, see (Doelling and Assaneo, 2021; Molinaro, 2025)).

Finally, our results also have important implications for the interpretation of multi-word synchronization findings. Apart from reflecting the processing of abstract linguistic structures (Ding, 2025, 2023; Ding et al., 2016), such findings have alternatively been explained in terms of single-word lexical processing (Frank and Christiansen, 2018; Frank and Yang, 2018; Kalenkovich et al., 2022) and implicit prosody imposed at phrase boundaries (Glushko et al., 2022). We observed striking functional and hemispheric differences between word-level (Figures S4 and S5) and phrase-/sentence-level IEPC (Figures 3 and 4) patterns, suggesting that multi-word processing cannot be reduced to single-word processing. The attenuated phrase-level responses with isochronous syllables/words also seem inconsistent with lexical accounts as isochrony should presumably increase the phase consistency of evoked responses. Accordingly, isochronous speech might have obscured the contribution of single-word lexical processing in previous frequency-tagging studies (Burroughs et al., 2021; Frank and Yang, 2018; Kalenkovich et al., 2022; Kazanina and Tavano, 2023; Lo et al., 2022; Zhao et al., 2024). The stimulus-independent responses in left STG (especially for sentences) are also difficult to explain in terms of implicit prosody, which nevertheless could be related to the stimulus-sensitive modulations of IFG (especially for phrases). Likewise, the lack of identical patterns between phrase- and sentence-level synchronization also supports the involvement of segregated neural responses that cannot solely result from implicit prosody imposed at either phrase or sentence boundaries. Our findings thus align with the structural interpretation of multi-word synchronization (although not necessarily with their hierarchy assumption (Frank and Christiansen, 2018; Lo et al., 2023)) and extend them to more natural anisochronous speech contexts.

## Conclusion

We provide MEG evidence for the fundamental dissociation between the processing of multi-word linguistic structures (phrases and sentences) and syllables using a novel frequency-tagging paradigm based on more natural anisochronous continuous speech. Multi-word structure processing relies on endogenous inferences that remain robust to the anisochrony of the acoustic signal, while syllable processing is instead maximized by artificial isochrony reflecting exogenous responses that closely follow speech acoustics. This functional dissociation is also observed as hemispheric asymmetry, with stimulus-independent leftward lateralization for multi-word processing (particularly in STG) but stimulus-driven rightward lateralization (or bilaterality) for both syllable and acoustic processing. Our key findings thus advance a more realistic characterization of the flexible neural mechanisms that enable the efficient comprehension of natural speech.

## Methods

### Participants

Thirty native German speakers (21 female; mean age = 28.87 years, standard deviation = 3.99 years) participated in this study. All participants were right handed and reported no history of neurological, hearing, nor language disorders. All participants provided written consent before the experiment and received a monetary compensation of 12 € per hour. The study was approved by the Ethics committee of the University of Leipzig (application number: 060/17-ek) and was conducted according to the Declaration of Helsinki.

### Stimulus and study design

Participants listened to German sentences composed of 4 bi-syllabic words (Adjective/Possessive + Animate Noun + Verb + Inanimate Noun: e.g., *Meine Eltern lesen Bücher,* ‘My parents read books’). The first two words formed a Noun Phrase (i.e., the subject) and the last two words formed a Verb Phrase (i.e., the verb and direct object). A total of 200 unique sentences were created following these linguistic criteria. Lexical frequency (log10 lemma frequency from dlexDB database) of subject (animate) nouns and object (inanimate) nouns was not significantly different (mean subject: 3.05, SD = 0.82; mean object: 3.07, SD = 0.83; t(398) = −0.30, p = 0.76). We next concatenated these sentences into long sequences, resulting in 20 different 10-sentence trials. Only the first word (Adjective/Possessive) was repeated (2-3 times) across sentences but never within trials nor at the same ordinal position between trials. Each of these 10-sentence trials was converted into audio files using a speech synthesizer (Natural Language API by Google) that preserved natural (i.e., time-varying/anisochronous) word durations. Similarly to natural speech features, synthesized trials involved continuous speech streams without acoustic pauses between adjacent syllables/words. The resulting speech stimuli were then manipulated using the Praat Vocal Toolkit (Corretge, 2012). First, syllable boundaries (80 per trial) were manually annotated by a native German speaker. Second, based on these annotated syllable boundaries, we imposed the same duration over specific linguistic units of interest with increasing duration (syllables < 4-syllable phrases < 8-syllable sentences) by selectively lengthening or shortening their original durations: 0.25 s for syllables (i.e., 4 Hz), 1 s for phrases (i.e., 1 Hz), and 2 s for sentences (i.e., 0.5 Hz). Isochrony was imposed separately to syllables/phrases/sentences and thus shorter isochronous units automatically turned longer units into isochronous but also units without imposed isochrony maintained the original time-variable/anisochronous duration distributions. Following this procedure, we created four different experimental conditions containing different combinations of isochronous (i.e., fixed ‘+’) and/or anisochronous (i.e., variable ‘-’) durations across syllables/phrases/sentences. More concretely, in the ‘syl+phr+sen+’ condition, the three linguistic units were isochronous as in the conventional frequency-tagging paradigm. The rest of conditions containing anisochronous units represent a novel aspect of the current frequency-tagging paradigm. In the ‘syl-phr+sen+’ condition, syllables were anisochronous but both phrases and sentences were isochronous. In the ‘syl-phr-sen+’ condition, both syllables and phrases were anisochronous but sentences were isochronous. In the ‘syl-phr-sen-’ condition, all linguistic units were anisochronous but the total duration of the trial was matched to 20 s as in the other conditions. To confirm that our timing manipulation effectively modified the duration of each linguistic unit in the intended manner, syllable boundaries from the resulting audios were again annotated by a native speaker and then we calculated the duration distribution of each linguistic unit (see Figure S1). As expected, isochronous units were systematically clustered to their intended mean duration across conditions, while anisochronous units were consistently distributed over a range of variable durations around their corresponding mean duration. We also calculated the duration of bi-syllabic words, which displayed similar duration patterns as syllables. Each 20-s trial had a version in the four experimental conditions. Finally, prosodic cues from speech stimuli were further removed by flattening their pitch (120 Hz) and normalizing their amplitude (65 dB). Speech materials of the current study are available in an OSF repository (https://doi.org/10.17605/OSF.IO/WSX9V).

We then computed spectral analysis (power spectrum) of the speech envelope to assess how the isochrony manipulation changed the presence of periodic acoustic modulations. Specifically, for each 20-trial and condition, we extracted the Hilbert envelope of the broadband signal, calculated the frequency-domain decomposition via Fast Fourier Transform (FFT; frequency resolution: 1/20 = 0.05 Hz), and squared the resulting Fourier coefficients. The speech envelope power spectrum revealed the most salient acoustic modulations at 4 Hz corresponding to the mean frequency of syllables (Figure 1D). We first assessed the statistical significance of acoustic peaks at the frequency of each linguistic unit based on a signal-to-noise ratio (SNR) measure obtained by comparing the power values at the mean frequency of each linguistic unit by comparing it to the mean of its neighbouring frequencies (+/− 0.4 Hz; one-sided *t*-test, uncorrected) separately for each condition. For the syllable frequency, the SNR results showed that the 4-Hz peak was statistically significant in the ‘syl+phr+sent+’ condition (t(19) = 19.463, p < .001), ‘syl-phr+sent+’ condition (t(19) = 8.233, p < .001), ‘syl-phr-sent+’ condition (t(19) = 3.803, p < .001), but not in the ‘syl-phr-sent-’ condition (t(19) = 0.246, p > .05). For the word frequency, the 2-Hz peak was statistically significant in the ‘syl+phr+sent+’ condition (t(19) = 2.518, p < .05), ‘syl-phr+sent+’ condition (t(19) = 2.769, p < .01), ‘syl-phr-sent+’ condition (t(19) = 2.508, p < .05), but not in the ‘syl-phr-sent-’ condition (t(19) = −0.321, p > .05). For the phrase frequency, the 1-Hz peak was statistically significant only in the ‘syl+phr+sent+’ condition (t(19) = 2.316, p < .05), but not in the ‘syl-phr+sent+’ condition (t(19) = 1.133, p > .05), ‘syl-phr-sent+’ condition (t(19) = 1.698, p > .05), and ‘syl-phr-sent-’ condition (t(19) = 1.153, p > .05). For the sentence frequency, the 0.5 Hz was not statistically significant in any condition: ‘syl+phr+sent+’ condition (t(19) = 0.372, p > .05), ‘syl-phr+sent+’ condition (t(19) = 0.215, p > .05), ‘syl-phr-sent+’ condition (t(19) = −0.149, p > .05), and ‘syl-phr-sent-’ condition (t(19) = 0.282, p > .05). The SNR results thus indicate that both syllables and words display robust acoustic modulations in the speech signal, particularly with isochronous durations. Moreover, the SNR results also suggest that the speech signal contains acoustic modulations at the frequency of phrases in the ‘syl+phr+sent+’ condition, which might challenge the endogenous interpretation of phrase-level effects. To better understand how acoustic modulations in the speech signal vary across conditions, we also run linear models to assess the effect of condition separately at the frequency of each linguistic unit (bonferroni-corrected t-test for post-hoc contrasts). For the syllable frequency, the 4-Hz peak was significantly different across conditions (condition effect: (F(3,76) = 169.4, p < .001). More concretely, this 4-Hz peak was stronger for isochronous than anisochronous syllables (‘syl+phr+sent+’ vs. ‘syl-phr+sent+’: t(76) = 14.780, p < .001; ‘syl+phr+sent+’ vs. ‘syl-phr-sent+’: t(76) = 18.330, p < .001; ‘syl+phr+sent+’ vs. ‘syl-phr-sent-’: t(76) = 20.352, p < .001). Moreover, we generally observed progressively reduced 4-Hz power across conditions with additional anisochronous units (‘syl-phr+sent+’ vs. ‘syl-phr-sent+’: t(76) = 3.549, p < .01; ‘syl-phr+sent+’ vs. ‘syl-phr-sent-’: t(76) = 5.572, p < .01; ‘syl-phr-sent+’ vs. ‘syl-phr-sent-’: t(76) = 2.023, p > .05). In contrast to syllables, we did not observe significant differences across conditions at the frequency of the rest of linguistic units: 2-Hz word frequency (F(3,76) = 2.173, p > .05), 1-Hz phrase frequency (F(3,76) = 1.0555, p > .05), and 0.5-Hz sentence frequency (F(3,76) = 0.078, p > .05). These results thus suggest that multi-word linguistic structures (i.e., phrases and sentences) cannot be processed based on acoustic cues but rather on abstract inferences independently of their isochrony.

Speech materials were presented in four different blocks, with each block only comprising trials of the same experimental condition. Two additional blocks involving word-order manipulations were also included but are not reported in the current study as they address different theoretical questions. All blocks were presented randomly across participants. Each trial started with a white cross presented in the middle of the screen. Audio onset was preceded by a jitter (0.5-1 s) and audio offset was followed by 2.3 s of silence. Participants were instructed to fixate the cross at all times and to blink when the cross disappeared between trials. To make sure that participants were paying attention to the stimuli, a sentence-recall task was used at the end of each trial. After the silence, a sentence appeared on the screen with two answer (yes/no; counterbalanced) options at the bottom. Participants were instructed to report whether they had listened to that exact sentence or not during the immediately preceding audio. Answers were provided by pressing two buttons (left-right button position was counterbalanced across participants) and visual feedback (correct vs. incorrect answer) was always presented after each response. To reduce repetition effects across trials from different conditions, the visually presented sentence always targeted a distinct sentence across conditions. Sentences requiring a ‘no’ answer were created by replacing a single word (counterbalanced across trials) with a same-category word from another sentence from that trial (counterbalanced across conditions/blocks).

### Data acquisition

MEG data were recorded in a magnetically shielded room (AK3b, Vacuumschmelze, Hanau, Germany) using a 306-channel Neuromag Vectorview MEG (MEGIN, Helsinki, Finland). MEG data were recorded with a sampling rate of 1000 Hz and an online low-pass filter of 330 Hz (no online high-pass filter). Subjects were comfortably seated up-right with a fixed head position. Stimuli were delivered using Psychtoolbox (Kleiner et al., 2007) in Matlab. Acoustic stimuli were presented binaurally through air-conduction earplugs (ER3– 14A/B; Etymotic Research Inc.) connected via a 50 cm plastic tube to piezo phones (TIP-300, Nicolet Biomedical). Visual stimuli were back-projected on a semitransparent screen positioned ∼90 cm from each participant using a projector (Panasonic PT-D7700E, Matsushita Electric Industrial) with a refresh rate of 60 Hz.

### Data preprocessing

MEG data were first pre-processed using the Signal-Space-Separation (tSSS) method implemented in Maxfilter 2.2 (Elekta-Neuromag) to remove external magnetic noise and correct for head movements. Further pre-processing steps were implemented in MATLAB R2021A (The MathWorks) via the FieldTrip toolbox(Oostenveld et al., 2011). First, continuous MEG data were segmented into single trials (20 s preceded by 0.5 s and followed by 1.5 s, both periods without acoustic stimulation) and then low-pass filtered at 90 Hz. Next, heartbeat and ocular artifacts were manually identified using Independent Component Analysis (ICA) and then linearly subtracted (Infomax algorithm from Fieldtrip). MEG magnetometers (102 sensors) were then discarded and only planar gradiometers (102 pairs of sensors) were kept for subsequent steps. MEG data were finally downsampled to 100 Hz.

### Source localization

T1-weighted Magnetic Resonance Images (MRI) were obtained from each participant with a 3T MRI scanner (Magnetom Trio, Siemens AG). Individual MRI images were separated into scalp, skull, and brain components using the segmentation algorithm implemented in Fieldtrip (Oostenveld et al., 2011). We used a standard Montreal Neurological Institute (MNI) brain template available in SPM toolbox for two participants whose MRI was not available. Co-registration of the anatomical MRI images with MEG signal was performed using the manual co--registration tool available in Fieldtrip. The source space was defined as an homogeneous 3D grid with a 5-mm resolution. Single-trial MEG signals (planar gradiometers) were low-pass filtered at 8 Hz (filter order = 2; FIR two-pass forward and reverse filter) before downsampling to 100 Hz. Next, whole-brain cortical sources of single-trial MEG signals were estimated using LCMV beamformer (Van Veen et al., 1997). Separately for each trial, the forward model was computed using a single-sphere head model for three orthogonal source orientations and only the first 2 principal components with maximum variance were kept based on the singular value decomposition (SVD) factorization. Source-localized single-trial signals were estimated for 162 sub-regions from the Human Brainnetome Atlas (Fan et al., 2016).

In addition, we selected predefined regions of interest for the subsequent analyses. These regions included the left and right Inferior Frontal Gyrus (IFG: dorsal, ventral, and opercular portions of BA 44; rostral and caudal portions of BA 45; and Inferior Frontal Sulcus) as well as the left and right Superior Temporal Gyrus (STG: BA 41/42 of the auditory cortex; rostral and caudal portions of BA 22; medial and lateral portions of BA38). All analyses were performed separately for each of the sub-regions within left/right IFG and left/right STG regions.

### Behavioral task

Given the behavioural ‘yes/no’ response in the sentence-matching task after each trial, we calculated the mean accuracy of each participant separately for each condition.

### Inter-Trial Phase Coherence (ITPC)

To confirm the presence of neural synchronization to the frequency of linguistic units, we computed ITPC resolved in the frequency domain as common in previous frequency-tagging studies exclusively based on isochronous speech (Gui et al., 2020; Har-Shai Yahav and Zion Golumbic, 2021; Lu et al., 2023; Makov et al., 2017; Sokoliuk et al., 2021). Following previous studies, we first removed the first sentence of each trial (∼2 s) to avoid transient responses driven by acoustic onset. Next, we decomposed each trial (18 s) into the frequency domain via Fast Fourier Transform (FFT; frequency resolution: 1/18 = 0.055 Hz) separately for each region and component. We extracted the phase from the resulting Fourier coefficients and then computed ITPC in terms of non-uniform distribution of single-trial phase angles in the circular plane (expressed as z-values) from the Rayleigh test using the ‘circ_rtest’ function from the Circular Statistics toolbox (Berens, 2009) in Matlab. ITPC was computed using the Rayleigh test (as in (Har-Shai Yahav and Zion Golumbic, 2021)) for consistency with the main phase synchronization analysis (see below). This analysis was implemented separately for 162 sub-regions, 2 principal components, and for each frequency in the range comprising the frequencies of interest (∼ from 0.5 to 4 Hz). Z-values were further normalized by dividing by the mean of the +/− 5 surrounding frequency bins (corresponding to +/− 0.275 Hz). Finally, we averaged the normalized z-values between the 2 principal components and then across sub-regions from IFG and STG regions as well as left and right hemispheres.

### Inter-Event Phase Coherence (IEPC)

To quantify phase synchronization to the immediate occurrence of linguistic units, we computed IEPC resolved in both frequency and time domains. We further epoched each 20-s trial including its preceding 0.5 s and its subsequent 1.5 s (both comprising periods without acoustic stimulation) to avoid edge artifacts in the time-frequency analysis. Next, separately for 162 sub-regions and 2 principal components, we computed the time-frequency representation of each 22-s trial using Morlet wavelets (at each frequency of interest in steps of 0.045 Hz and 0.01 s) as implemented in Fieldtrip (ft_freqanalysis; cfg.method = ‘wavelet’; cfg.output = ‘fourier’; cfg.width = 3; cfg.gwidth = 3). For consistency across conditions with higher/lower isochrony, we computed the time-frequency analysis on the frequency ranges of the ‘syl-phr-sent-’ condition only comprising anisochronous units (syllables = 2.55-7.27 Hz; words = 1.55-2.77 Hz; phrases = 0.82-1.18 Hz; sentences = 0.45-0.55 Hz) for all conditions. We then extracted the instantaneous phase of the resulting Fourier coefficients at boundaries corresponding to the exact occurrence of each linguistic unit (i.e., offset of the preceding unit and onset of the following unit) based on manual annotations by a native German speaker. For phrase boundaries, we focused on the first phrase boundary within each sentence (between the noun phrase and the verb phrase) as the second phrase boundary (verb phrase) was confounded with the sentence boundary. We discarded the instantaneous phase from the first and last boundaries of each trial to avoid trial onset and offset effects, respectively. This resulted in a series of instantaneous phases concatenated across all 20 trials - separately for syllables (n = 1560), words (n = 760), phrases (n = 160), and sentences (n = 160) - and then IEPC was computed using the non-uniformity’s Rayleigh test as in the previous ITPC analysis with the ‘circ_rtest’ function from the Circular Statistics toolbox (Berens, 2009) in Matlab. Z-values from the 2 components were summed and then further normalized by dividing by the mean computed across 162 sub-regions and 4 conditions. Finally, z-values were averaged within each sub-region from IFG and STG regions separately for the left and right hemispheres.

### Lateralization Index (LI)

To assess IEPC hemispheric asymmetry across conditions, we computed a LI as LI = right/(left + right) for each linguistic unit. This LI was calculated following (Brodbeck et al., 2022), in our case using the normalized IEPC separately for 162 (i.e., 81 bilateral) sub-regions and then averaged within each sub-region from IFG and STG regions.

### Temporal Response Function (TRF) of speech envelope

To examine the processing of speech acoustics in the time domain, we model the linear relationship between neural responses and the continuous speech envelope of our stimuli by means of ridge regression following a forward (encoding) Temporal Response Function (TRF) approach (Crosse et al., 2021, 2016). Specifically, the forward TRF approach can be described as a linear model that estimates a TRF filter weight that optimally maps stimulus features (e.g., speech envelope) to neural responses. The TRF filter weight was estimated by minimizing the deviance (mean squared error) between empirical and predicted neural responses using the regularized (ridge regression) solution. To avoid overfitting, we tested a range of regularization terms [lambda values: (10^−2^ - 10^5^)] using a ‘leave-one-out’ cross-validation procedure by maximizing the Pearson’s correlation between the observed and predicted responses.

The TRF analysis was implemented using the functions from the mTRF toolbox (Crosse et al., 2016) in Matlab. Specifically, *mTRFcrossval* was used for the cross-validated selection of lambda values, *mTRFtrain* for the estimation of the filter weight, and *mTRFpredict* to test the estimated model. The speech envelope was computed using *mTRFenvelope* (downsampled to 100 Hz; root-mean-square compression to log10(2)) after applying a zero-phase low-pass filter at 8 Hz (filter order = 2; IIR two-pass forward and reverse filter) and normalized to root mean square of all trials across conditions. For consistency with the acoustic signal, neural responses were further low-pass filtered at 8 Hz following the initial source-localization procedure (see above) and then normalized by dividing by the standard deviation of all trials across conditions and sub-regions. We used Pearson’s r to assess the goodness of fit between the empirical and predicted neural responses of the left-out trial. To avoid overfitting, we first concatenated every two trials (resulting in 10 40-s trials) so that training and test partitions corresponded to 80% and 20% of the data, respectively. To focus on post-stimulus neural responses, we selected positive time lags ranging from 0 to 0.2 s. This procedure was computed separately for each participant, sub-region (n = 162) and principal component (n = 2). Finally, the r values from the 2 principal components were summed and then averaged across trials. Given that the speech signal contained acoustic correlates for syllables and that syllable-level IEPC was strongest in STG, we focused this analysis only on left and right STG regions.

### Statistical analysis

Statistical significance at the group level was determined with linear models in RStudio. Specifically, we use the function *lm* to model the effect of independent variable(s) on each dependent variable and assess its statistical significance using the function *anova* together with *emmeans* for pairwise comparisons (bonferroni corrected two-sided *t*-test). Independent variables corresponded to condition (‘syl+phr+sent+’, ‘syl-phr+sent+’, ‘syl-phr-sent+’, ‘syl-phr-sent-’), region (IFG, STG) and hemisphere (left, right). We use the following linear models for different dependent variables: *behavioral accuracy ∼ condition*; *ITPC (for each frequency) ∼ condition*; *IEPC (for each frequency) ∼ condition*region*hemisphere*; *TRF prediction accuracy (Pearson’s r) ∼ condition*hemisphere*. In addition, following the spectral analyses of the speech envelope, statistical significance of ITPC was initially tested with a SNR by comparing the ITPC of each frequency to the mean of the +/− 4 surrounding frequency bins (corresponding to +/− 0.22 Hz; one-sided *t*-test). For the lateralization analysis, LI values were statistically compared against the empirical value representing perfect bilateral distribution (i.e., 0.5; *t*-test, two-tailed, corrected for multiple comparisons).

## Data and code availability

The pre-processed data and code supporting the findings of this study will be made accessible after publication.

## Acknowledgements

We thank Antonia Schmidt, Laura Riedel, Paula Baer, and Johanness Gereons for help with stimulus preparation, Burkhard Maess for support in experiment setup, and Yvonne Wolff-Rosie for assistance in data collection.

## Author contributions

conceptualization: J.M., N.M., L.M.

methodology: J.M., G.D.L., L.M.,

formal analysis: J.M.

writing – original draft: J.M., N.M., L.M.

writing – review & editing: J.M., G.D.L., N.M., L.M.

investigation, data curation, visualization: J.M.

funding acquisition: N.M., L.M.

supervision: N.M., L.M.

## Supplementary materials

**Figure S1.**
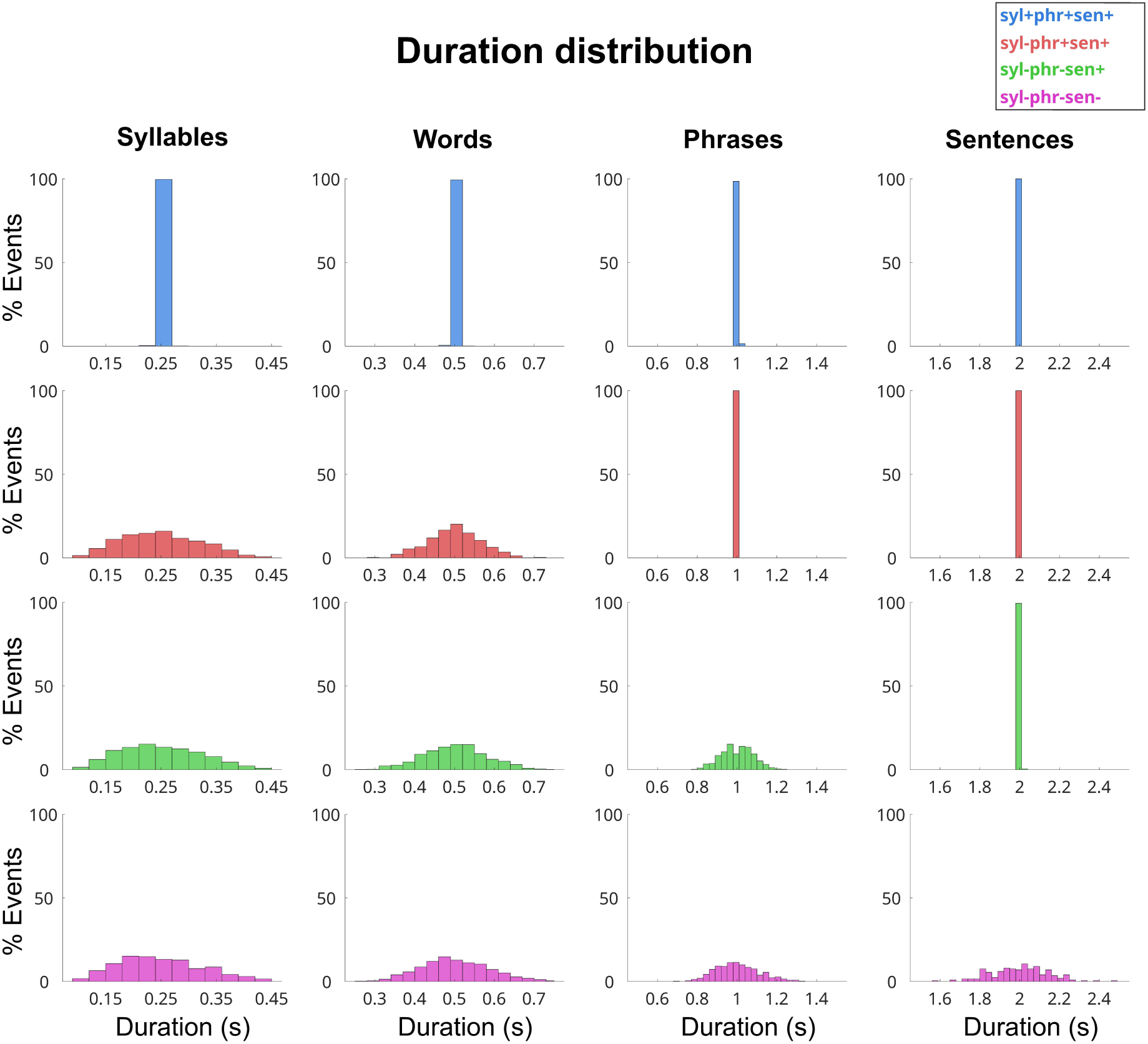
Duration distribution of linguistic units in continuous speech stimuli. Histograms showing the percentage (%) of units (X-axis) as a function of their duration (Y-axis; in seconds, s) derived from the manually annotated stimuli, show separately across conditions representing the isochronous-to-anisochronous continuum (in rows; color-coded; see Legend) and each linguistic unit of interest (in columns; here including words). This shows the efficacy of our intended duration manipulation: isochronous (“+”) units are clustered to a single duration but anisochronous (“-”) units are distributed over a range of durations.

**Figure S2.**
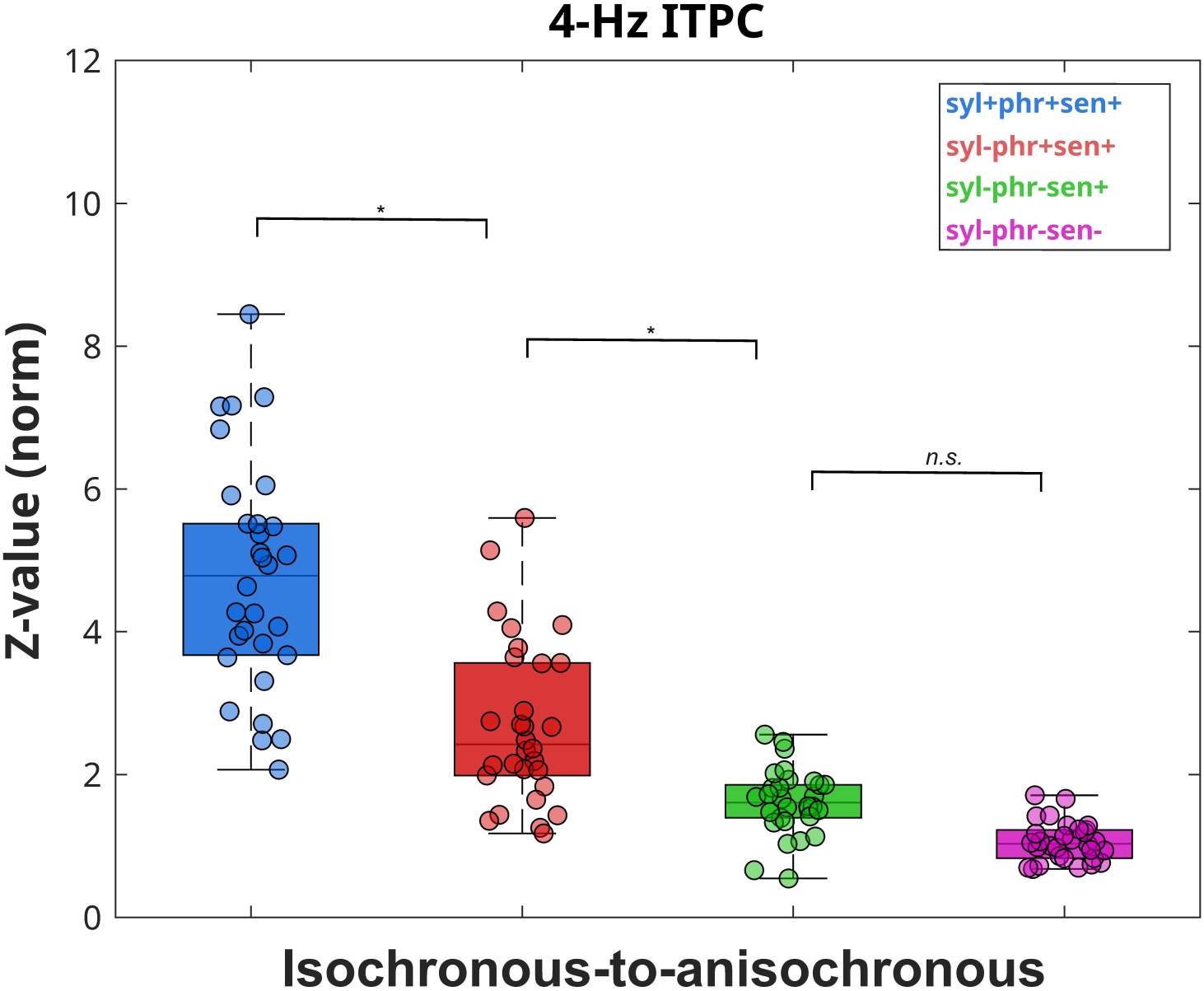
ITPC comparison across conditions at 4-Hz syllable frequency. The 4-Hz ITPC peak was significantly different across conditions (condition effect: F(3,116) = 77.765, p < .001). The condition with isochronous syllables resulted in significantly stronger 4-Hz ITPC compared to the rest of conditions with anisochronous syllables (‘syl+phr+sent+’ vs. ‘syl-phr+sent+’: t(116) = 7.844, p < .001; ‘syl+phr+sent+’ vs. ‘syl-phr-sent+’: t(116) = 12.003, p < .001; ‘syl+phr+sent+’ vs. ‘syl-phr-sent-’: t(116) = 14.130, p < .001). Mirroring spectral analysis on the speech envelope power spectrum, this 4-Hz ITPC peak was progressively reduced across conditions containing additional anisochronous units (‘syl-phr+sent+’ vs. ‘syl-phr-sent+’: t(116) = 4.159, p < .001; ‘syl-phr+sent+’ vs. ‘syl-phr-sent-’: t(116) = 6.286, p < .001; ‘syl-phr-sent+’ vs. ‘syl-phr-sent-’: t(116) = 2.127, p > .05).

**Figure S3.**
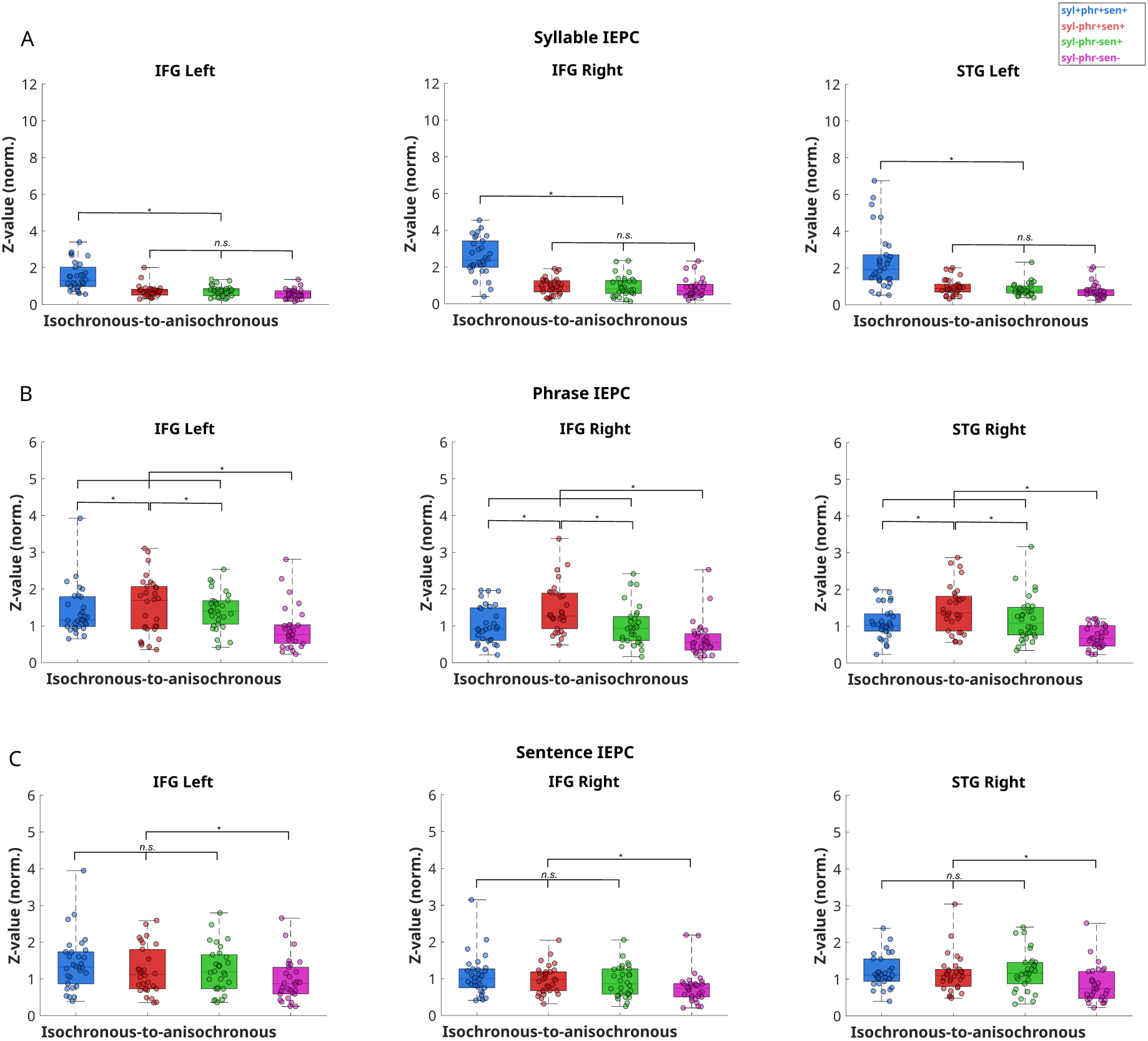
Inter-Event Phase Coherence (IEPC) secondary results (see Figure 3 for details). IEPC for the regions with the less strong effects (regions with the strongest effect in Figure 3) for syllables (A), phrases (B), and sentences (C). Asterisks represent statistically significant differences between conditions.

**Figure S4.**
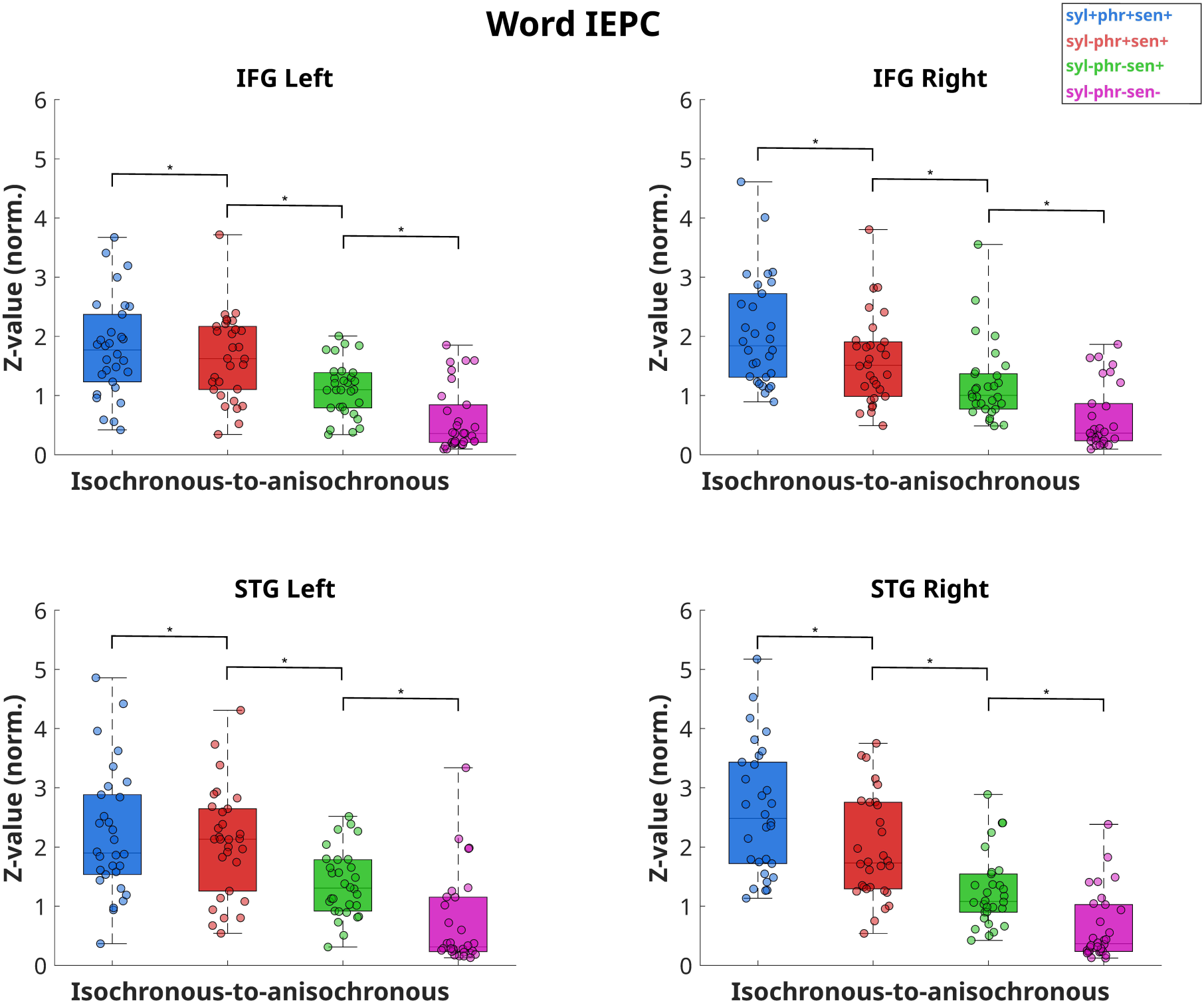
Inter-Event Phase Coherence (IEPC) for word boundaries (see Figure 3 for details). Word-level IEPC for all regions. Asterisks represent statistically significant differences between conditions. **I**EPC at word boundaries was significantly different across conditions (condition effect: F(3,464) = 92.274, p < .001), showing significantly higher IEPC for isochronous compared to anisochronous words (‘syl+phr+sent+’ vs. ‘syl-phr+sent+’: t(464) = 3.590, p < .01; ‘syl+phr+sent+’ vs. ‘syl-phr-sent+’: t(464) = 9.471, p < .001; ‘syl+phr+sent+’ vs. ‘syl-phr-sent-’: t(464) = 15.470, p < .001). Similarly to the speech envelope power and ITPC spectra, word-level IEPC was significantly lower across conditions progressively containing additional anisochronous units (‘syl-phr+sent+’ vs. ‘syl-phr-sent+’: t(464) = 5.881, p < .001; ‘syl-phr+sent+’ vs. ‘syl-phr-sent-’: t(464) = 11.881, p < .001; ‘syl-phr-sent+’ vs. ‘syl-phr-sent-’: t(464) = 6.000, p < .001). Word-level IEPC was significantly higher in STG than in IFG (region effect: F(1,464) = 16.342, p < .001) but not significantly different between hemispheres (hemisphere effect: F(1,464) = 0.561, p > .05; region x hemisphere interaction: F(1,464) = 0.198, p > .05) consistently across conditions (condition x hemisphere interaction: F(3,464) = 1.666, p > .05; condition x region interaction: F(3,464) = 1.954, p > .05; condition x hemisphere x region interaction: F(3,464) = 0.261, p > .05). This suggests bilaterality independent of isochrony.

**Figure S5.**
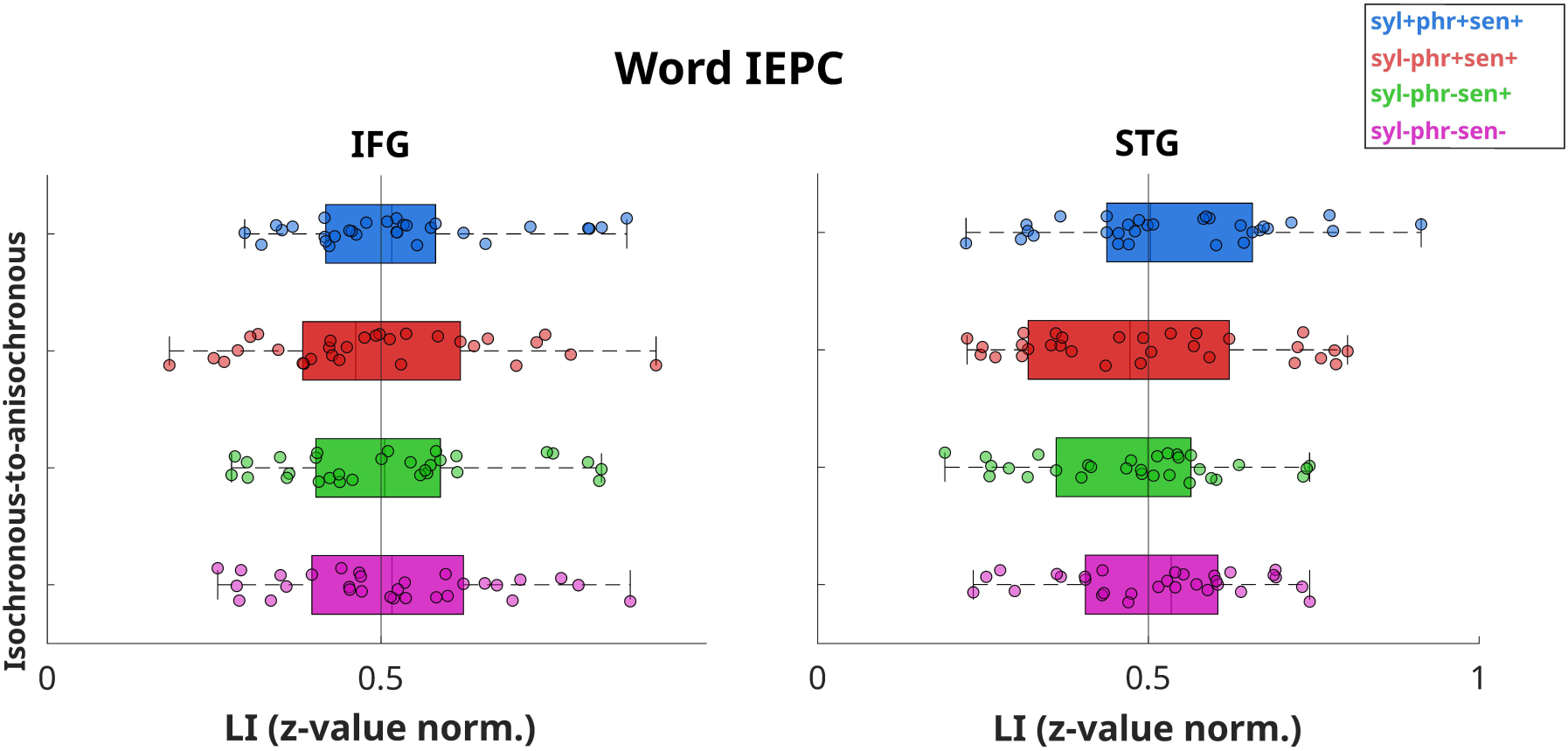
Lateralization Index (LI) of Inter-Event Phase Coherence (IEPC) results for words (see Figure 4 for details). We did not observe significant word-level IEPC lateralization in any condition for neither IFG nor STG, reinforcing the previous IEPC results that suggested bilateral distribution independent of isochrony.

**Figure S6.**
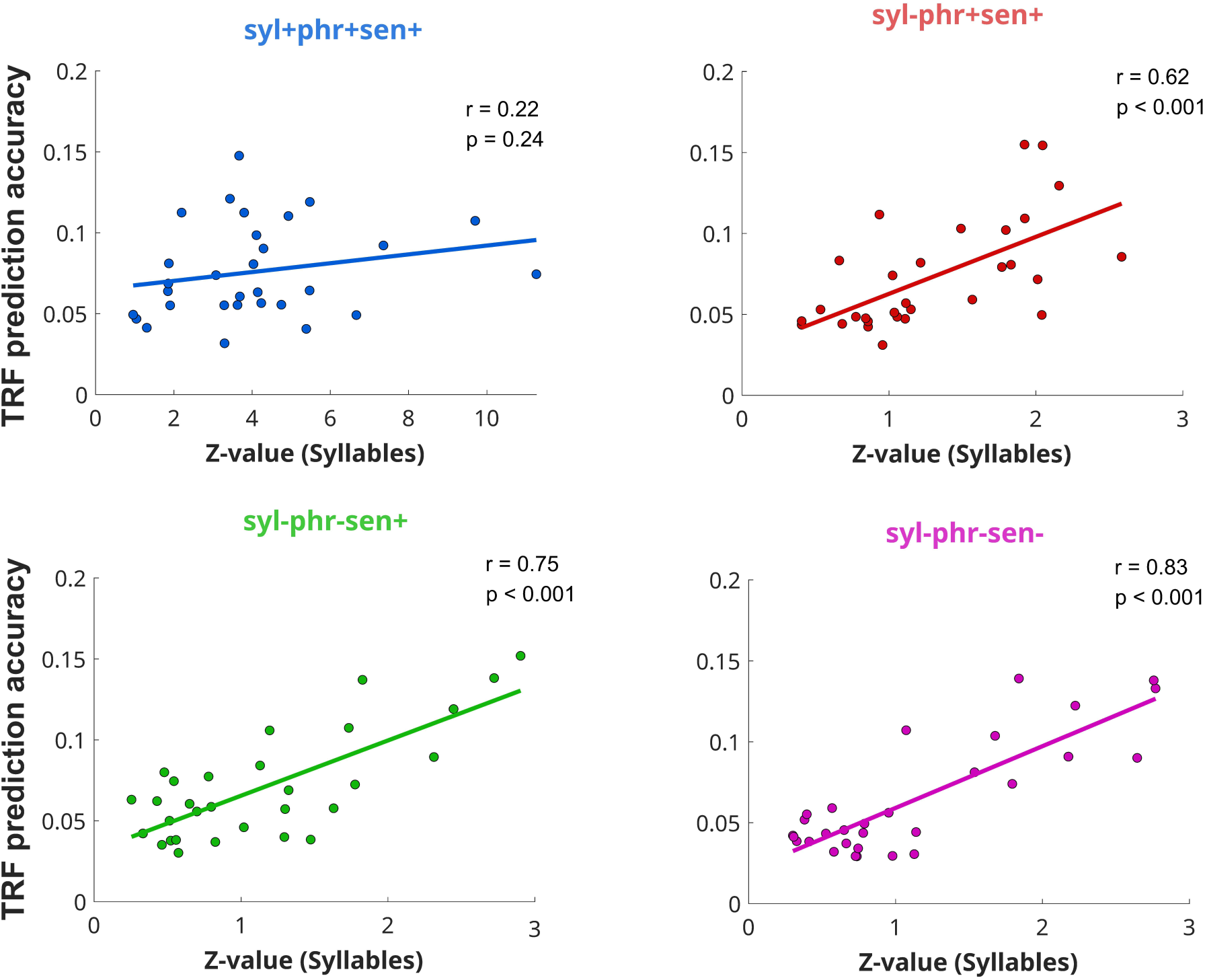
Correlation between Inter-Event Phase Coherence (IEPC) and Temporal Response Function (TRF) results. Correlation (Pearson’s r) at right STG obtained between syllable-level IEPC (Figure 3A; X-axis) and TRF prediction accuracy (Figure 5B; Y-axis), shown separately for each condition (color-coded) along with their corresponding correlation coefficient (r) and statistical significance p-value (p; uncorrected).

